# ARID1B damaging variants from more than one million genomes, cause human diseases by impairing protein-protein interactions, stability, and regulation

**DOI:** 10.64898/2026.06.19.733400

**Authors:** Jessica B. Wagenknecht, Neshatul Haque, Jane Gresser, Xiaowei Dong, Michael T. Zimmermann

## Abstract

*ARID1B* is the most frequently *de novo* altered gene across a spectrum of human neurodevelopmental disorders and cancers. We found 1,456 missense variants in ARID1B’s C-terminal domains, 94% of which are clinically uninterpreted and 59% of which are observed in human disorders and cancers. Using integrative modeling of ARID1B cBAF and DNA bound structures, we calculate how these variants impact key functional sites, structural stability, and topological features, finding that 617 variants clearly impair ARID1B’s interactions within BAF, interactions with DNA, fold stability, or post-translational regulation. Our study enables scalable and precise variant interpretation by illuminating the molecular mechanisms by which missense variants damage ARID1B function, with commonalities across malignancies and congenital anomalies mutually informing therapeutic developments.

## Introduction

Population scale genetics sequencing aims to improve health for all. To meet that vision, new approaches are needed to interpret the myriad ultra-rare genetic variations that are being discovered. Neurodevelopmental disorders (NDDs) are a heterogeneous collection of underlying genetic conditions including Coffin-Siris syndrome (CSS), autism spectrum, and non-syndromic intellectual disability, potentially affecting as much as 7% of the population and representing a substantial fraction of germline rare diseases ^1,2^. AT-rich interaction domain 1B (*ARID1B*) is among the most frequently mutated genes in NDDs ^3^. Consulting diverse genomic resources, we have identified that almost every variant observed in *ARID1B* (97%) is ultra-rare or unique to individuals, making statistical approaches unviable on a per-variant basis. We expect that the less severely affected side of the spectrum, and extent of allele penetrance, is incompletely characterized. At the same time, ARID somatic alterations are frequent in human tumors. Further, there is a strong reciprocal relationship between germline and somatic diseases with respect to ARID genes - *ARID1A* is 2x to 4.5x more frequently altered in human tumors than *ARID1B* ^4,5^, while *ARID1B* variants are considerably more common in NDDs than *ARID1A* ^3,6^. Consequently, pharmacology is rapidly progressing in the druggability of cancers with altered ARID-family genes by leveraging the commonality of *ARID1A* somatic loss-of-function that creates a tumor vulnerability to ARID1B pharmacologic degraders ^7–10^. Therefore, it is critical to the administration of new therapies, to understand better which of the many rare missense variations are functional. All told, the clinical actionability for ARID1B variation is increasing, motivating the current study to identify mechanistic subgroups among the broad genomic landscape of this essential gene.

The ARID1B protein (also known as BAF250b) binds other proteins, chiefly SMARC-family proteins like SMARCA4, to form a BRG1/BRM-associated factor (BAF) complex. BAF is an epigenetic regulator which remodels nucleosome positioning to alter gene expression ^11,12^. Different combinations of proteins form different BAF complexes with specific physiologic and cellular functions (e.g., cBAF, ncBAF, and pBAF) ^13^. ARID proteins play a critical role within BAF complexes by connecting the core module proteins to the ATPase module proteins via interactions of armadillo (ARM) repeats ^14,15^. All ARID proteins also share an AT-rich interaction (ARID) domain (for which they were named) that binds to DNA. An ARID1 Scaffolding Domain 1 (ASD1), with homology to Drosophila’s Eld/Osa, ^16^ is also present, which appears to support BAF complex assembly ^9^. Thus, the function of ARID1B variation interpreted according to its role in the BAF complex and chromatin regulation will be superior to the gene investigated in isolation.

In the current study, we use large scale human cohorts that cumulatively and conservatively comprise over a million samples, to map somatic and germline variation across ARID1B. Our long-term goal is to gain insight into how germline and somatic variation in ARID1B shapes human development, brain function, and cancer. Further, we seek to resolve how ARID1B gene dosage, protein function, and context determine disease mechanisms. To start addressing these goals, we first reclassify all 1,456 distinct missense variations observed in the combined cohort, 94% of which are currently VUS, according to how they affect intra- and inter-molecular properties. We find five distinct mechanistic classes which further illuminate which variants are damaging and how, to the normal functions of ARID1B within canonical BAF complexes. Then, we quantify how different functional classes of germline variants stratify the combined cohort according to observed phenotypes and cancer tissue-specific patterns. We find most pathogenic variants are destabilizing, and most variants we identify as functionally-damaging are associated with germline or somatic diseases. In this way, we add to the global characterization of ARID1B variation using deep structure-based analysis.

## Results

### Characterizing the Broad ARID1B Genomic Landscape Across Human Conditions

To characterize the ARID1B genomic landscape, we first aggregated human genetic variation from multiple large-scale sources spanning population studies, to congenital genomics databases with phenotype and inheritance data, and of variants found in human tumor samples. Unifying across these six databases (AllofUs, gnomAD, COSMIC, CBioPortal, LOVD, and ClinVar; see **Methods**), we identified 13,028 unique DNA variations in the *ARID1B* gene region. Most of these variants (7,111; 55%) have only been observed in population studies. A smaller number of variants are unique to congenital (1,156; 9%) or cancer genomics (2,482; 19%), with a few (429; 3%) found within all three (**Figure 2**). Population studies recapitulate 578 cancer-associated variants, but over twice as many disease-associated variants (1,088) are also found in population studies. This data suggests that there may be additional functional variants within the population which have not yet been characterized, leaving wide the possibility even for deep phenotyping of carriers as candidates for understanding better the genotype-phenotype associations for this pleiotropic gene. The overlap between cancer and congenital disease databases was even smaller (184; 1%). Cancer-associated variants are more unique from population-level variants than heritable disease-associated variants are. Finally, there is high overlap between both population databases (**Figure 2B**) and comparatively less overlap between either the two congenital or the two cancer databases (**Figure 2C, 2D**), suggesting that there may be many disease-associated variants not yet catalogued, which will also require means of specific and mechanistic interpretation such as we develop and use herein.

**Figure 1:**
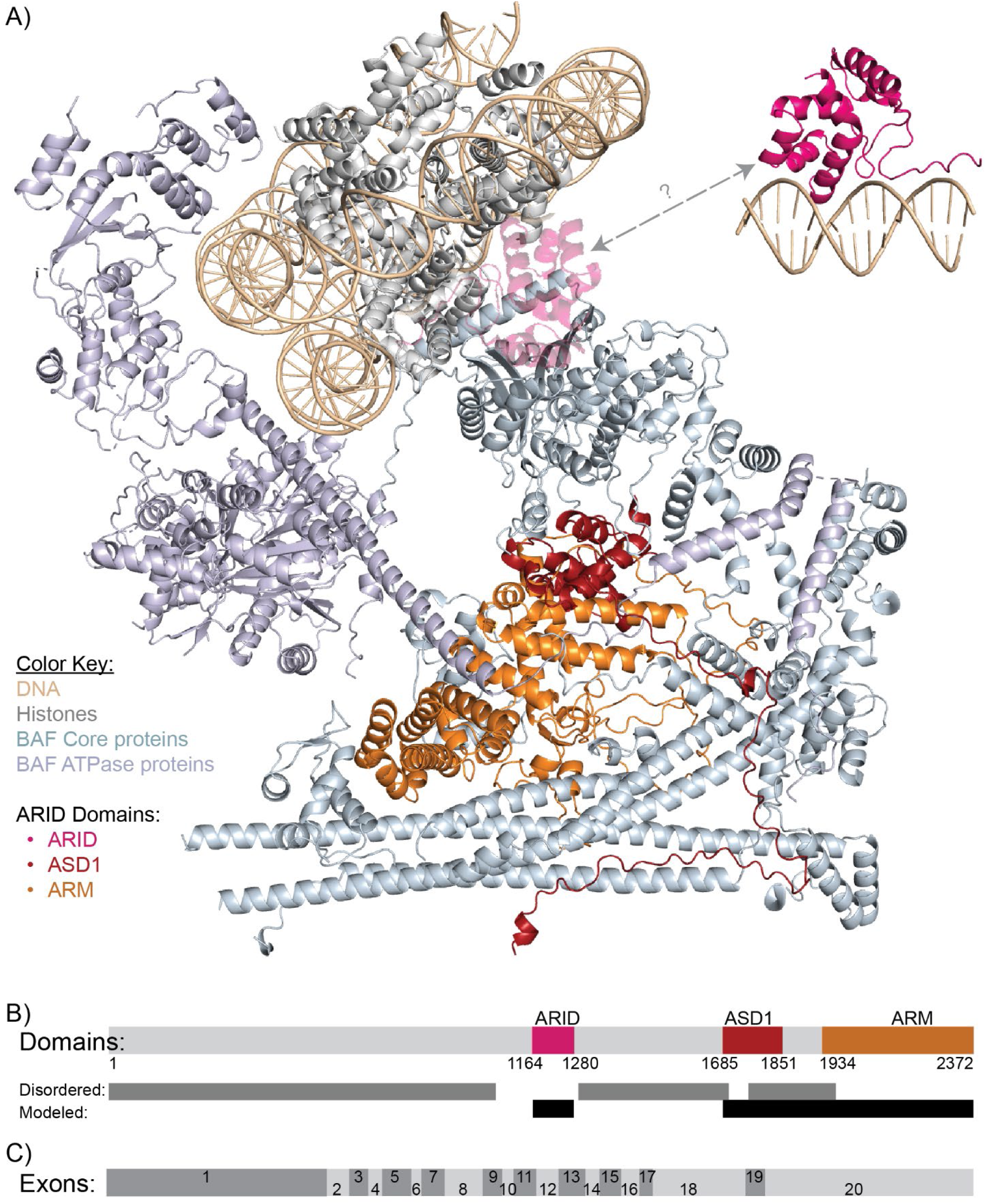
The ARID protein is central to the BAF complex, connecting the core proteins and ATPase proteins. The core proteins here (SMARCA, …) are colored light blue, and ATPase proteins (SMARCA4, …) are colored light purple. Histone proteins are shown in grey and the DNA in tan. Finally, ARID1B’s three structured domains are colored independently: the ARID domain is bright pink, ASD1 domain is dark red, and ARM domain is orange.

**Figure 2:**
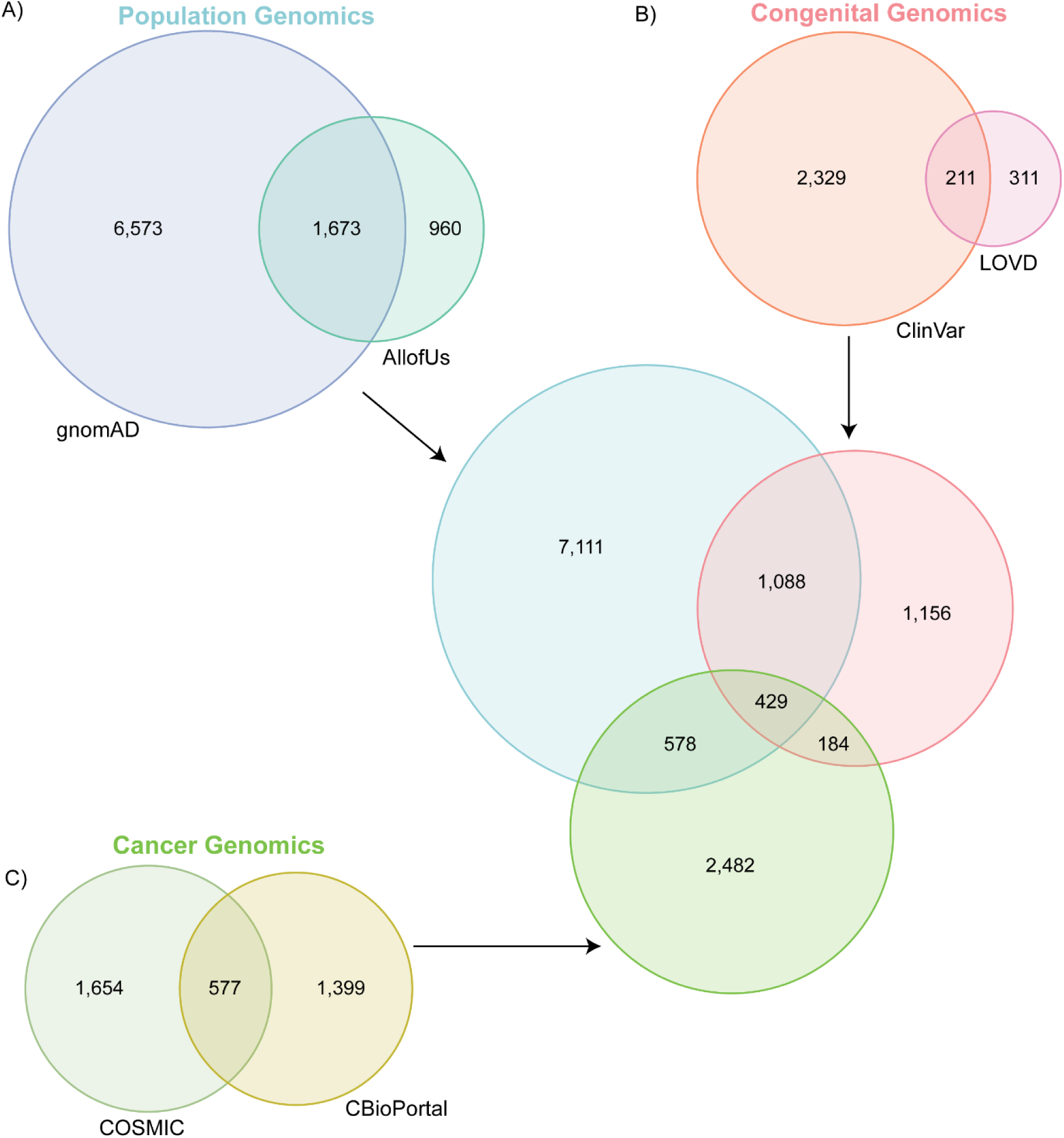
Thousands of variants are observed in ARID1B, most being uniquely observed by clinical ascertainment context. a) a Euler diagram of ARID1B variants from population genomics databases gnomAD and AllofUs. B) a Euler diagram of mendelian disease -associated ARID1B variants from databases ClinVar and gnomAD. C) a Euler diagram of ARID1B variants found in tumor samples from databases COSMIC and CBioPortal. The variants within these three genomics types come together to form a Euler diagram of the population genomics, mendelian disease-associated, and tumor-specific ARID1B variants.

### ARID1B Alterations are associated with a large range of congenital phenotypes and malignancies

Since *ARID1B* is associated with intellectual and developmental disabilities, we investigated the pathogenicity of variants, and their associated phenotype or trait spectrum. We found that more variants are VUS (1,058) than benign (977) or pathogenic (820). Yet, the majority (77%) of the VUS are missense variants, while most pathogenic variants (90%) are pLoF, and many benign variants (60%) are silent (**Table 1**). Further, 41 variants had interpretations in one database but were VUS in another. Most variants (82%) are of unknown inheritance (or one or more of patients with that variant had unknown inheritance), and 158 variants are only found *de novo*. These *de novo* variants are almost entirely absent from population databases (96%), pathogenic (97%), and with high impact (84%). While less than half of the variants (46% of ClinVar; 44% of LOVD) had any phenotype or diagnosis annotations, the most common diagnosis is, as expected, CSS (602 variants), followed by inborn genetic diseases (572 variants), generic NDDs or intellectual disability (276), and ARID1B-related disorder (265). Looking beyond syndromic annotations, we find that the most common granular traits are various facial dysmorphologies (129) – including coarse facial features (33), thick eyebrow (27), and abnormal facial shape (26) – followed by growth impairments (114), vision or eye symptoms (70), altered digits (64), and hair phenotypes (64) – all hallmarks of CSS. Then, we investigated the co-occurrence between phenotypes, finding the more common phenotypes are also highly correlated (**Figure 3A**). More interestingly though, we found that heart symptoms (e.g., defects or abnormalities of heart physiology) are correlated with feeding difficulties and scoliosis, which are less common CSS symptoms and may represent a more mildly (compared to CSS) affected disease subgroup. Delayed speech or language is highly associated with delayed motor skills, suggesting a shared molecular cause underlying both; delayed motor skills is also associated with joint hypermobility. Seizure, abnormalities within the corpus collosum, and behavioral symptoms are also correlated, hinting to shared alterations to the brain. Overall, *ARID1B* alterations are associated with a broad spectrum of phenotypes across various body systems.

**Figure 3:**
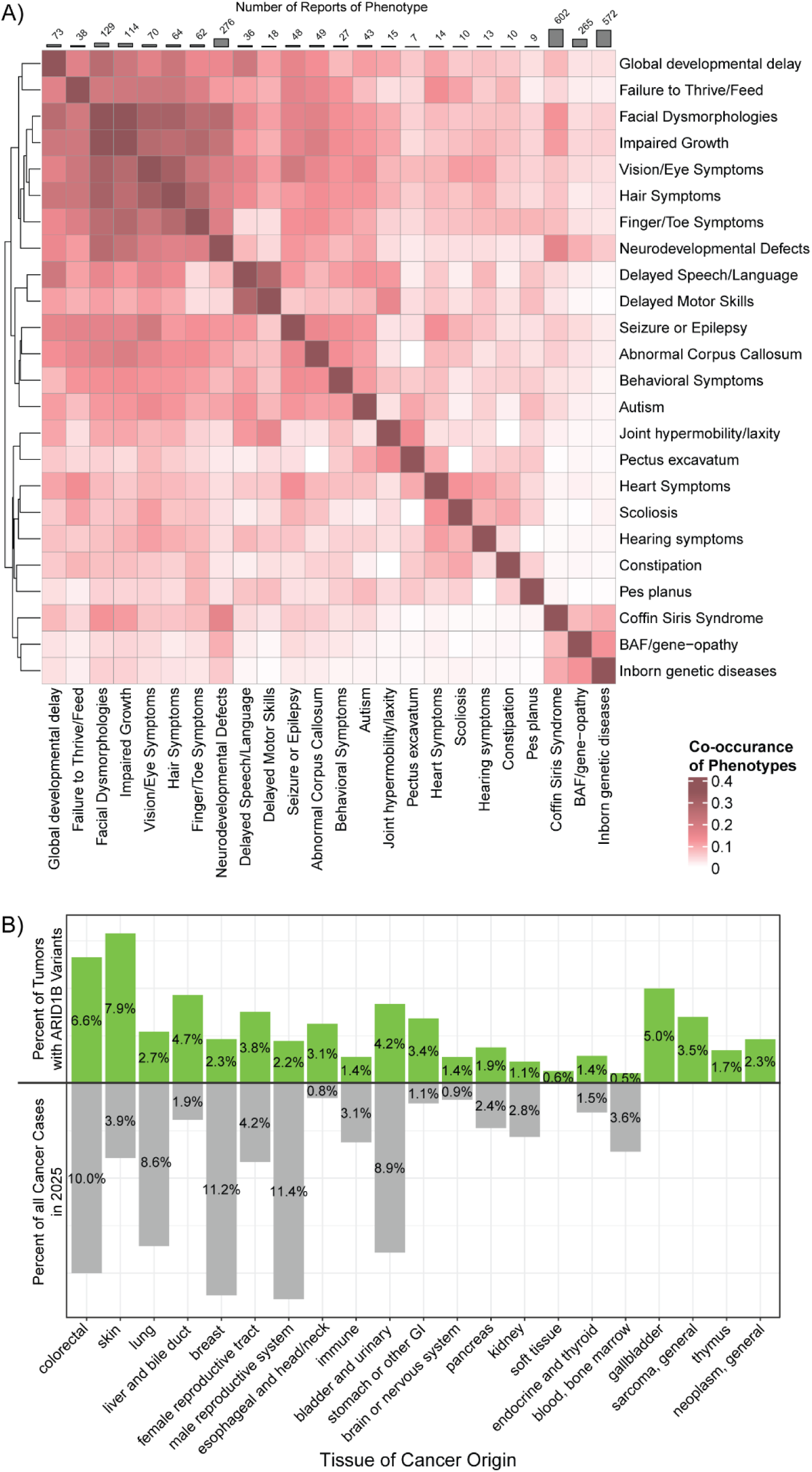
Variants in ARID1B are associated with a range of traits and cancers. A) the ratio of variants associated with the same trait divided by the overall number of variants gives the co-occurrence of traits, revealing that the more classical CS symptoms are commonly found together but some pockets of less-common traits co-associate. B) the percent of tumor samples in that tissue with an ARID1B variant is shown ordered by most to least total ATID1B variants; on the negative Y axis, the percent of new cancer cases in that tissue in 2025 is shown ^42^.

**Table 1:** Variant interpretation is highly correlated to impact level. , where pathogenic variants are primarily high-impact, benign primarily low impact, and VUS primarily moderate.

|  |  | Variant Impact Level |  |  |  |
| --- | --- | --- | --- | --- | --- |
|  |  | High | Moderate | Low | Modifier |
|  |  | *Frameshift, start loss, stop gain, or splice acceptor or donor loss* | *Missense or inframe insertion/deletion* | *Splice region variant or synonymous* | *UTR or intronic variant* |
| Clinical Significance | Pathogenic | 702 | 96 | 12 | 10 |
|  | VUS | 41 | 942 | 49 | 26 |
|  | Benign | 1 | 376 | 483 | 117 |
|  | Not in Clinical Databases | 555 | 4289 | 1353 | 2617 |

Next, we investigated the cancers associated with *ARID1B* variation. Overall, most cancer-associated variants are of moderate impact (missense or inframe insertions/deletions); however, liver and bile duct, pancreas, and kidney cancers have a far larger proportion of variants (≥50%) which are modifiers (i.e., intronic or in UTRs). The most common tissue in which *ARID1B* variant-tumors were found was colorectal (1,067 instances), followed by skin (693) and lung (569). However, colorectal, skin, and lung cancers were also within the top ten most samples tested. Hence, tissues with the highest percentage of *ARID1B* variants are skin (7.9%) and colorectal (6.6%), followed by gallbladder (5%) and liver or bile duct (4.7%) (**Figure 3B**). When compared to how common these cancers are overall, this is surprising; while colorectal cancers are fourth most common, liver and gallbladder cancers are rare. On the other hand, more prevalent cancers such as prostate and breast had a small percentage of *ARID1B* variants. This demonstrates a unique bias by tissue of origin, highlighting a potentially significant role of ARID1B in these tissues.

Population studies contained the largest number of variants in *ARID1B*, 79% of which are yet unobserved in human diseases. These variants, however, are remarkably individualized: 97% are ultra-rare, with almost half of those (4,027) only observed in one person each. On the other hand, only 89 variants have a minor allele frequency (MAF) greater than 1×10^-3^. This skew towards ultra-rare and person-specific variants demonstrates the need for large-scale research of ARID1B to better analyze variants using biologic and biophysical mechanisms, not only statistical associations, so that milder yet clinically important subgroups can be identified.

### ARID1B alterations differ regionally in type, frequency, and patient impact

Next, we explored the distribution of congenital, cancer, and population variants across the length of the protein (**Figure 4**). Given that prior clinical studies have revealed that many inherited and variably expressed variants in ARID1B are in exon 1 (spanning amino acids 1-597 in the RefSeq transcript) ^17,18^, we expected to find a different pattern in this region. Indeed, our study has revealed that exon 1 not only contains the most distinct variants (3,376), but also is especially rich in high allele-frequency variants (p = 2.2×10^-16^, Log_2_(odds ratio) = LOR = 1.01) and depleted for cancerous variants (p = 2.2×10^-16^, LOR = −2.06), when compared to the rest of the protein. This is an intrinsically disordered region (IDR), with no experimental structure and an amino acid composition that strongly indicates that it will not form secondary structure elements.

**Figure 4:**
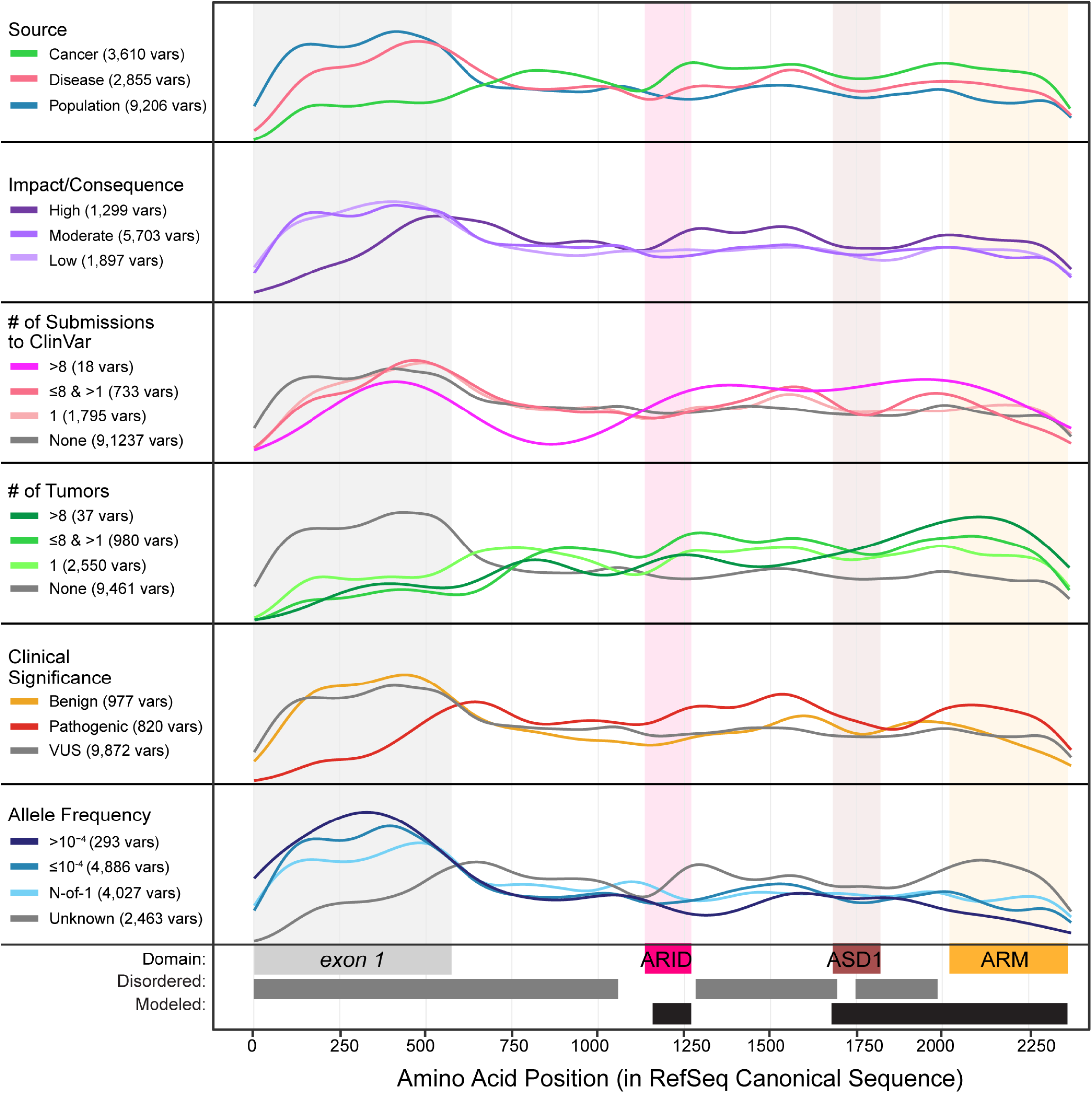
Five regions of ARID1B are enriched for different types of variants. The distribution of variants of specific categories across the protein sequence is shown, with domains and key regions shown, as well as existing experimental structures.

Interestingly, this region is depleted for high impact variants (p = 2.2×10^-16^, LOR = −1.15) or *de novo* variants (p = 1.4×10^-9^, LOR = −2.12). Patients with variants in this region are more often given the non-specific label “inborn genetic diseases” (p = 8.357×10^-7^, LOR = 1.01) than patients with variants elsewhere in the protein (**Supplementary Figure 1**). Further, exon 1 is significantly depleted in variants associated with facial dysmorphologies (p = 5.7×10^-5^, LOR = −1.79), impaired growth (p = 4.0×10^-5^, LOR = −2.06), and neurodevelopmental disorders (p = 1.2×10^-7^, LOR = −1.64). Thus, genetic patterns strongly point to this N-terminal disordered region as functionally distinct from the domains that participate within the BAF core particle.

Next, we wanted to determine if other regions of the protein had distinct patterns in variant associations. We found that all IDR regions except for those encoded by exon 1 (residues 598-1163, 1281-1684, and 1852-1933) are enriched for cancer occurrence (p = 2.2×10^-16^, LOR = 0.75). The second IDR, between the ARID and ASD1 domains (residues 1281-1684), also contains more high-impact variants (p = 1.3×10^-4^, LOR = 0.49). On the other hand, structured regions (defined as residues 1164-1280, 1754-1851, 1934-1948, 2056-2106, and 2133-2375) have more variants absent from the population (p = 2.2×10^-16^, LOR = 1.10). The ARM domain also has more pathogenic variants (p = 4.1×10^-5^, LOR = 0.63), demonstrating that the structured regions are already more well-characterized than disordered regions. Overall, the differences in variant distributions across the protein reveal distinct regions in ARID1B, especially demonstrated in the concentration of cancer-associated variants in disordered regions. Each of these distinct sections will require specific approaches to investigate, and we will start in this manuscript with investigating the structured domains.

### Missense variants impact ARID1B stability, regulation, and BAF formation

Most high and low-impact variants seen clinically are interpreted (93%), but most VUS are missense (89%). Thus, we focused on the conspicuous need for interpretation in ARID1B’s missense variants. In total, there are 4,946 unique protein alterations in ARID1B (see **Methods**), with only 56 currently classified as pathogenic and 251 as benign. To analyze how missense variants impact ARID1B functioning, we modeled the ARID domain bound to DNA, and the ASD1 and ARM domains of ARID1B within the cBAF complex (see **Methods**), creating two novel models of ARID1B. Within these modeled regions, 1,456 missense variants successfully mapped: 194 in ARID, 293 in ASD1, 811 in the ARM, and 158 variants between the ASD1 and ARM domains. Twice as many variants in these domains are associated with cancer (651) as mendelian diseases (315), in part because a higher proportion of cancer-associated variants were missense compared to those associated with congenital conditions.

We analyzed these variants for their potential impact on key known functions of ARID1B. Namely, 1) loss of BAF core assembly, 2) loss of BAF ATPase assembly, 3) loss of DNA interactions in the ARID domain, 4) loss of structural stability of ARID1B, and 5) loss of PTMs and motifs that may regulate BAF activities or localizations. We discovered 617 variants have impaired these ARID1B functions (**Figure 5A**). First, 186 variants lead to loss of (or damage to) BAF core assembly. In 71 of these variants, this effect is due to loss of key interactions between ARID1B and a BAF core protein (specifically, SMARCD1, SMARCC2, SMARCE1, or SMARCB1). For example, R2344W is ultra-rare and has been observed in CS syndrome and a grade III astrocytoma tumor. Our study reveals that R2344W has critical interactions both with itself and with SMARCD1, especially a hydrogen bond to SMARCD1 E170 (**Figure 6A**); it is also highly destabilizing, with a ΔΔG_folding_ = 5.9 kcal/mol and ΔFI = 9.7. The other 115 variants cause BAF core assembly loss by destabilizing ARID1B very near to these interaction sites. For example, E2333K has been observed in 36 tumors (33 esophagus tumors and 3 colorectal adenocarcinoma tumors) and is ultra-rare. Our study reveals that E2333K is highly destabilizing (ΔFI = 3.571, loses internal bonds) and is adjacent to SMARCB1 in the BAF core; a loss of stability there would likely impair SMARCB1 binding stability as well (**Figure 6B**). Second, 92 variants damage BAF ATPase assembly: 28 variants through loss of critical interactions to SMARCA4, and 64 through destabilization of ARID1B:SMARCA4 interaction regions. For example, D1788Y is associated with three breast cancer tumors and has never been seen in the population. Our study reveals that D1788Y is destabilizing (ΔΔG_folding_ = 3.3 kcal/mol) and causes a loss of a salt bridge between ARID1B and SMARCA4 R447 (**Figure 6C**). Third, 28 variants damage ARID domain binding with DNA: 2 variants through loss of interactions with DNA, and 26 through destabilizing the ARID domain adjacent to DNA. For example, R1246C is an ultra-rare variant associated with uterine endometrial cancer. We found that this variant is frustrating (ΔFI = 3.03) and causes a loss of a salt bridge to the phosphate backbone of DNA (**Figure 6D**). Fourth, 244 variants lead to loss of stability of ARID1B as a whole, which may still make it less capable of sustaining BAF protein assembly. Finally, 44 variants may alter the regulation of ARID1B: 10 by losing specific PTMs, and 34 by losing predicted regulatory motifs. Partitioning genetic variation into these five groups based on molecular function enables focused follow up study to deconvolute the associations among rare variation and each mechanism.

**Figure 5:**
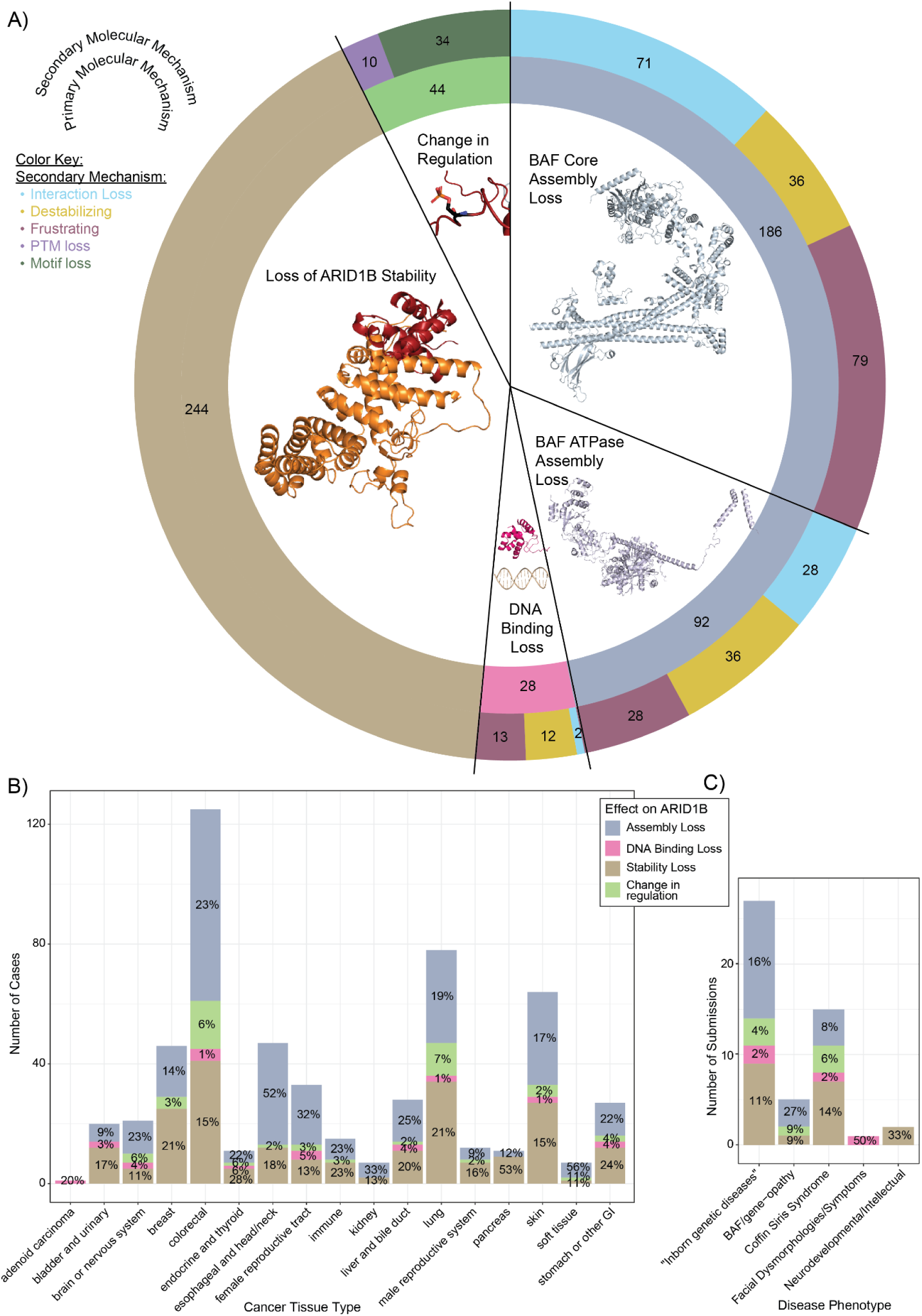
Variants in ARID1B cause loss of protein-protein interaction or destabilization of ARID1B. A) The number of variants which damage ARID1B are shown by the molecular mechanism by which they do so, and further broken down into their secondary mechanisms. B) The number of tumor samples in COSMIC, ClinVar, and CBioPortal with variants damaging ARID1B by these mechanisms are shown by the main tissue group where the tumor was found. The percent of all cases for that tissue type is labeled. C) Similarly, the number of submissions to ClinVar or LOVD are shown by mechanism and associated phenotype.

**Figure 6:**
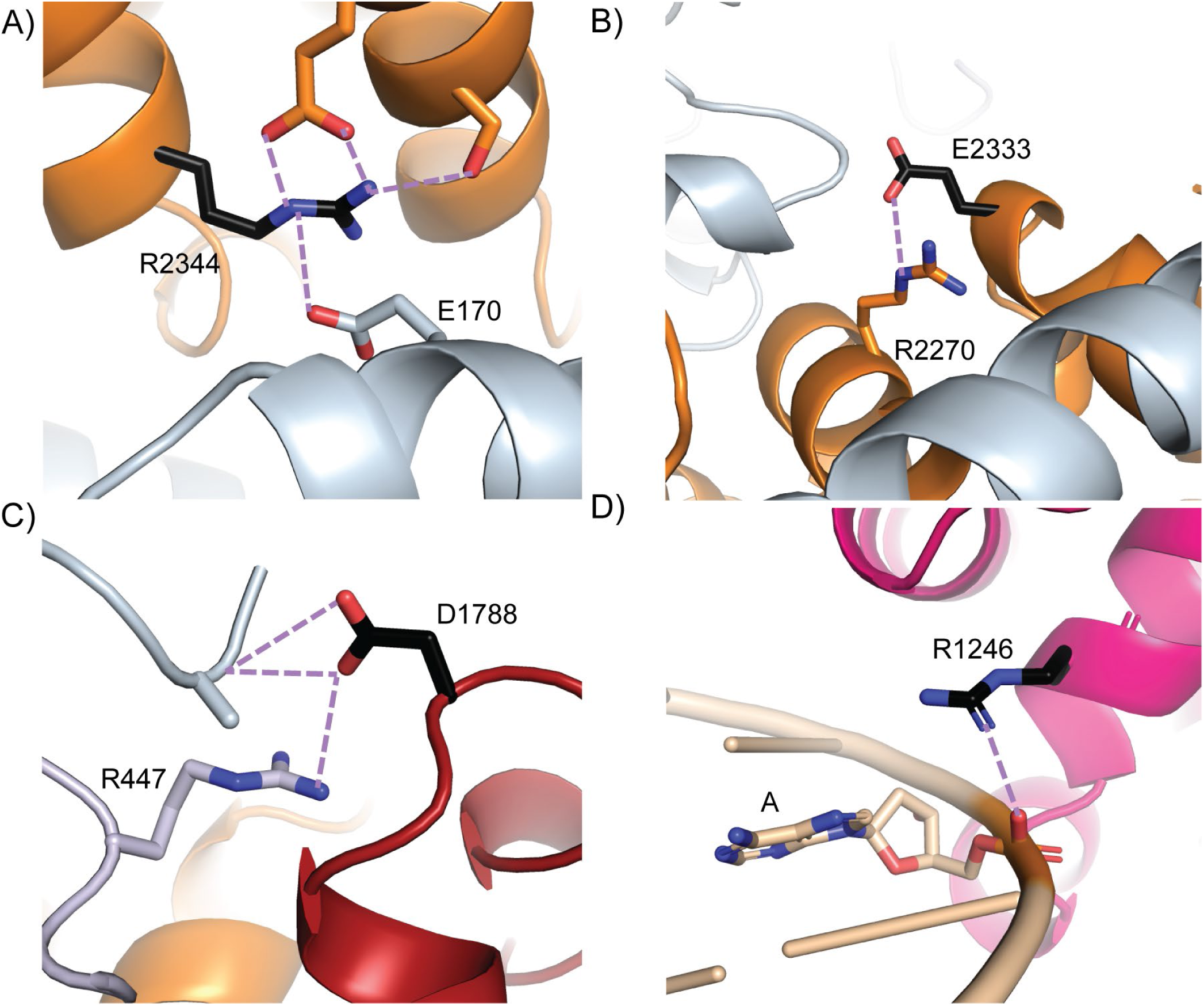
Examples of variants which damage ARID1B functions. A) R2344W (shown in black) impairs SMARCD1 binding (blue) via loss of hydrogen bond to SMARCD1 E170. B) E2333K impairs SMARCB1 binding due to loss of stability adjacent to SMARCB1 binding. C) D1788Y impairs SMARCA4 binding due to loss of a salt bridge between ARID1B D1788 and SMARCA4 R447. D) R1246C loses interactions with DNA due to loss of a salt bridge to the phosphate backbone.

Not only does this analysis provide mechanistic characterization of variants effects, it also reveals which variants damage ARID1B function. This functional information can improve variant interpretation as an enduring resource and reference point for future variant discovery. Throughout the structured and disordered regions in the domains we studied, we found 589/1,372 VUS (43%) and 14/24 pathogenic (58%) have evidence of potential damaged function (**Supplementary Figure 2**). However, we also found a potential mechanism of effect for 14 of the 60 benign variants (23%) in structured regions, suggesting that not all may be entirely tolerated by the protein. For example, L2302V (MAF = 2.9×10-5) is considered benign but has been submitted to ClinVar 4 times for CSS, ARID1B-related disorder, and inborn genetic diseases. We found that it is energetically frustrating and loses an internal hydrogen bond, destabilizing ARID1B domain structure.

Similarly, A2129V is considered benign but has been observed in three tumors (intrahepatic cholangiocarcinoma, soft tissue sarcoma, and anaplastic carcinoma) and submitted to ClinVar. We found that it loses a critical hydrogen bond to SMARCD1 protein, impairing BAF core assembly. Therefore, our analysis also delineates which variants in ARID1B are damaging, including hundreds of VUS but also a handful of benign variants which may not be benign in all contexts or for all patients.

Next, we analyzed whether any cancers or phenotypes were more associated with one mechanism over others. We found that overall, cancer-associated variants were mildly enriched for having a mechanistic explanation (p = 0.019, LOR = 0.37), with 298 of the 617 damaging variants being associated with cancer (46%), compared to 39% of congenital disease variants. Not only are there more missense variants associated with cancer than congenital disease (as stated above), but more of the cancer-associated variants damage ARID1B functions. We did not find these damaging variants enriched for any specific tissue (**Figure 5B**), and no specific phenotypes were enriched for damaging variants (**Figure 5C**), however, only 106 variants in structured regions had specific phenotypic data to begin with, so the power to detect statistically significant associations is low. Pathogenic variants are more likely to have evidence of damage (p = 0.0041, LOR = 2.17) and are more often associated with stability loss (p = 0.012, LOR = 1.61) (**Supplemental Figure 2**). Overall, our analysis reveals the mechanisms by which 387 variants may contribute to tumorigenicity or rare disease, applicable to the 721 people behind the submissions to CBioPortal, COSMIC, ClinVar, or LOVD.

Our identification of underlying molecular mechanisms enables the opportunity to extend the interpretation of existing experimental data. For instance, the impact of missense ARID1B ARM domain variants was recently measured with respect to protein abundance and cellular lethality ^19^. Between the two studies, there are 826 variants shared; within these, we explain the specific mechanism behind 76 of the 79 variants which had decreased abundance (**Supplemental Figure 3**), with 75/76 affecting stability in one way or another, among other impacts. Likewise, we find that the 6 variants with the lowest proliferation in the ARID1A^-/-^ assay are highly destabilizing (ΔΔG_folding_ > 3 kcal/mol) and/or frustrating (ΔFI > 4), and they all cause loss of an internal hydrogen bond. However, we also find 269 additional variants that may impact stability with smaller or more localized effects as to not outright diminish ARID1B abundance. We also study an additional 630 variants beyond the regions they studied and find another 217 destabilizing variants and six regulation-impacting variants. Therefore, we both explain and expand upon the results of this assay.

## Discussion

This study followed from our previous experience in rare disease genetics whereby we have identified that mechanistic structure-based analyses can clarify the functional significance of numerous variations and including those private to individuals ^20,21^. Importantly, through this methodology, underlying mechanisms are revealed for the dispositioned and VUS alike. By jointly querying more than 1.4 million samples spanning diverse clinical and population settings, we gathered the largest-to-date compendium of human variation in ARID1B. This transcription factor is a critical component of the chromatin remodeling complex BAF, and we identified loss of BAF assembly as a consistent theme for its dysregulation across congenital and somatic conditions. Further, our analyses indicate that many currently uninterpreted variants may damage ARID1B functioning.

While ARID1B is regarded as a missense-tolerant gene (Z = −2.94) ^22^, we studied the missense variation because we found that within healthy adults, the majority of variation (especially polymorphic variation) is within the low-complexity disordered region exclusively encoded by the first exon (**Figure 3**), while missense variation within structured domains have been associated with disease and shown to cause loss of function in cells ^19,23–25^. Thus, we anticipated that polymorphic variation in ARID1B has different physiology from missense substitutions in the ordered domains and the gene is in fact intolerant to variation of the latter type.

Therefore, we focused on the structured components of ARID1B that form the central organizing point of the BAF core particle. As such, our study expands on prior studies of the impacts of missense variants on ARID1B functioning, finding the same variants destabilizing ^19^ as measured by decreases in cellular protein content – we have expanded our search to investigate additional mechanisms of damaging protein-protein interactions and protein-DNA interactions, and have investigated variation across the all structural domains. Our results also align with the study of E. Bosch et. al, which found six destabilizing missense variants in ARID1B lead to protein aggregation rather than degradation ^26^ – variants we also find destabilizing to the BAF complex. Thus, integrated analyses that account for multiple lines of evidence yield the most sensitive characterization of ARID1B genetic variation.

Future studies of the intrinsically disordered regions and non-missense variants would shed even further light on functional versus non-functional variation in ARID1B. While CSS and ARID1B-related disorder are most commonly caused by haploinsufficiency, work on ARID1B and related BAF components also discusses dosage-dependent effects in terms of copy-number changes, partial deletions, mosaicism, or hypomorphic missense variants ^27–29^. Further, noncoding regulatory alterations of ARID1B are only just beginning to be characterized. For example, a recent study identified a conserved ultra-conserved element (UCE_11311) acting as an enhancer of ARID1B, with noncoding mutations that alter ARID1B expression ^30^. When we mapped variants onto the 3D structure, we discovered that many of the benign variants within the ARM domain are in disordered loops, while fewer pathogenic variants are in loops. That there are pathogenic variants in these loops highlights the specificity needed to accurately predict effects of single ultra-rare variations, and additional functional significance which may be overlooked by the lack of secondary structure elements. Even in long intrinsically disordered regions lie established pathogenic and potentially function-altering variants. Specifically, variants in exon 1 have been noted for causing milder phenotypes and occasionally being inherited from seemingly unaffected parents ^17,18^. We found that variants in this region were especially associated with disease, but were more often low or medium impact variants than in other regions (**Figure 4**). Interestingly, this region also contains most of the population variation. What precisely causes this first exon to be so different from the rest of the disordered region, its specific functions, and how variants may cause their different phenotypes, should be further studied. One current hypotheses is that an alternative translation start site may cause the variable expression ^18^. Yet, functional studies have shown a high functional significance for the N-terminal IDR in ARID1B overall, and regions in exon 1 control localization and expression of other genes ^31^. Though our analysis greatly extends the number of variants with in-depth study, further analysis on this region will complement the current study.

We also identified that patient-derived *de novo* variants are almost entirely absent from population databases (96%) and are pathogenic (97%) with high impact (84%). Potentially, this is due to genetic testing patterns in rare disease populations and the inheritance pattern alone better fitting paradigms for interpretation. This reveals a gap in precision medicine – not all patients will receive as in-depth of study of variant effects (inheritance testing, in vitro study, etc.), and those without this attention are less likely to receive a diagnosis, especially with low or medium -impact variants. While it is not realistic for all variants to be investigated *in vivo* or *in vitro*, all variants can be investigated *in silico*, bringing more insights to all patients and increasing equity in precision medicine. The current study demonstrates that structural calculations can yield significant insights beyond what is naturally possible with sequence-based predictors. While many practitioners remain appropriately cautious about inferences derived from computational models, the rapid expansion of AI is reshaping expectations for variant interpretation. In parallel, we are rigorously evaluating and deploying computational structural genomics ^32–35^. We anticipate that the readiness of the genomics field for understanding, leveraging, and applying computational structural genomics to diagnostics, and beyond, is increasing. Such advances are important for answering critical questions such as: do germline ARID1B variants contribute to inherited cancer risk, either directly or via interaction with tissue-specific oncogenic processes? What is the underlying pathobiological mechanism that can link cancer risk, ASD contributions, and CS causality? Further, our mechanistic hypotheses for variant effects can be validated with further *in vitro* studies, with assays to examine BAF assembly and ARID1B stability. As Mermet-Meillon, et al. has already demonstrated the significance of destabilizing mutations in targeting ARID1B cancers with protein-based degraders, our concordant results inform the design of anti-BAF therapies.

In conclusion, we have analyzed the associations of 13,028 variants across *ARID1B* with respect to mendelian disease phenotypes, malignant tumors, and population frequency, finding notable co-occurrences of phenotypes and ARID1B-specific cancer tissues. Further, we analyzed 4,946 missense variants for mechanistic evidence of a damaging effect to the ARID1B protein, finding cause for concern in 617 variants, 95% of which are currently uninterpreted. Variants impair BAF assembly and structural stability of ARID1B, and may alter regulatory mechanism through sequence motifs and PTMs. These mechanistic characterizations rely on our novel integrated structure of ARID1B within cBAF, and not only distinguish damaging from potentially tolerated variants, but also provide rationale for variant effects – rationale which can be tested or can serve in future studies to advance treatments for cancer and mendelian disease patients alike.

## Methods

### Collection of sequence-based data

Genomic variants from the entire ARID1B genomic region were retrieved from the following databases at the given dates: from ClinVar on 1/28/2026 ^36^, from LOVD’s ARID1B database (curated by Gijs Santen) on 1/6/2026 ^37^, from AllofUs on 2/26/2026 ^38^, from gnomAD v4 on 10/20/2025 ^22^, from COSMIC’s variants database on 4/1/2026 ^39^, and from CBioPortal’s curated set of non-redundant studies (see supplementary information for complete list of which studies) on 1/21/2026 ^40^. Individual-level phenotype information from the ARID1B LOVD database was also gathered. As variants from LOVD and CBioPortal were provided in GRCh37 format, we converted them all to VCF format using Ensembl’s GRCh37 variant recoder tool ^41^, then used Ensembl’s assembly converter tool to convert to GRCh38. Then, the HGVS formatted variants from all databases were combined to one dataframe and submitted to Ensembl’s VEP to retrieve isoform-specific annotation and HGVSp. In this process, 82 fusion variants from CBioPortal were filtered as they have no HGVS format, 8 variants from LOVD were filtered as they had too long of insertions or deletions to be submitted to Ensembl (over 64,000 bases), and another 66 complicated indels. Further, data from the initial 244,908 nucleotide bases were not present in COSMIC because transcript ENST00000635849.1 was the longest up-to-date transcript offered by COSMIC, but the RefSeq canonical transcript is longer. Variants in both AllofUs and gnomAD had similar allele frequencies (R^2^ = 0.94). For comparison to overall cancer rates, we relied on the 2025 study by the American Cancer Society ^42^.

For our protein-level analysis, we selected the RefSeq canonical transcript (NM_001374828.1). Importantly, 161 additional variants produce missense changes in other transcripts, but not in the RefSeq canonical transcript. However, only 5 of those variants are adjacent to structured regions, so by using the longer RefSeq transcript, our structure-based analysis captures almost all known structural regions – only transcript ENST00000635928.1 includes 21 additional amino acids at the end of the ARM domain.

Domains were defined as the following using the RefSeq Sequence: ARID as residues 1164-1280, ASD1 as 1685-1851, and ARM as 1934-2372. MobiDB was cross-referenced for evidence-based definitions of disordered regions, ignoring regions smaller than 50 aa ^43^. We analyzed the linear sequence motifs for potential regulatory interactions using ELM ^44^, and any with a probability of randomly occurring greater than 0.5%, or not within the nucleus, were filtered out. Post-translational modifications were identified by PhosphSitePlus ^45^. Finally, we obtained the properties of each reference and alternative amino acid using the IMGT amino acid properties ^46^.

### Collection of structure-based data

A 3D structure model of ARID1B’s ARID domain bound to DNA was created using Alphafold3 ^47^ and compared to the existing x-ray crystal structure (PDB ID 2EH9) ^48^, which aligned with RMSD = 0.443. A structure model of ARID1B’s ASD1 and ARM domains within the BAF complex was generated using Alphafold3, with the following sequences for structured portions of proteins ARID1B may interact with: ARID1B residues 1685-2372, SMARCA4 residues 401-470, SMARCD1 residues all 121-500, SMARCC2 residues 451-950 (twice), SMARCE1 residues 175-285, and SMARCB1 residues 1-385. For visualization purposes, this structure was aligned with the nucleosome-bound cBAF structure (PDB ID 6LTJ) to include the nucleosome and other BAF components ^15,49^.

These structure models were used to calculate the impact of variants on ARID1B. First, interactions between ARID1B sidechains and itself, other BAF proteins, and DNA were predicted using PLIP ^50^. Then, the distances between ARID1B residue C_α_ and other proteins’ C_α_ was calculated using Bio3D’s distance matrix ^51^. Other proteins were considered nearby when both C_α_ were within 10 Å. Then, FoldX was used to generate structures of each missense variant within ARID1B in the cBAF model described above (variants in the unstructured loops were excluded from this analysis – this includes amino acids 1691-1753, 1852-1933, 1949-2055, and 2107-2132) ^52^. Finally, the impact of these variants on structural stability was calculated by comparing the FoldX total folding energy for the variant and WT structure (giving ΔΔG_folding_), and the FrustratometeR frustration index (FI) of conformational, mutational, and single residue decoys ^53^. ΔFI levels ≥ 1 were considered frustrating, and ΔΔG_folding_ > 1.8 was considered destabilizing.

### Data Integration for Variant Analysis

First, we determined whether variants altered any of the annotations, including motifs, PTMs, and interactions. Variants’ impacts on motifs were determined using the regular expressions provided by ELM. We generated the altered sequence within the same region the motif was found previously, and tested whether the regular expression still found the motif after alteration. When a variant was at the exact site of a PTM, alteration of that PTM was determined by testing whether the new amino acid was still useable for that specific modification. For that, we considered S, T, and Y viable for Phosphorylations, R viable for methylarginine, and K viable for acetylated lysine, SUMOylated lysine, and ubiquitinated lysine. When a variant was within a phosphorylation motif, the regular expression was used as described for other motifs.

Finally, when a variant was within 3 amino acids of a phosphorylation and its change in volume was greater than 40 Å^3^ or it flipped charges, we considered it possibly altering to that phosphorylation due to altering the local environment.

Then, we determined whether variants altered interactions using the change in amino acid properties and the current interaction. In detail, residues which formed a salt bridge, but the variant lost its charge, were considered to lose their salt bridge interaction. Variants at residues where sidechains donated electrons for a hydrogen bond, but changed to a non-donating residue, were considered to lose that interaction, and vice-versa for hydrogen-accepting residues. Variants which lost interactions to SMARCD1, SMARCC2, SMARCE1, or SMARCB1 were considered to impact protein-protein interactions to BAF core proteins, while variants which lost interactions to SMARCA4 were considered to impact protein-protein interactions to BAF ATPase proteins. Variants which lost interactions with other ARID1B residues were considered to damage ARID1B stability.

After summarizing every variant’s potential impact on motifs, PTMs, interactions, and stability, as well as their domain and proximity to interacting proteins/DNA, we summarized the implications of each variant by their most significant impact. Variants which lost specific protein-protein or protein-DNA interactions, or lost stability and were nearby other proteins, were summarized as damaging BAF assembly; variants which lost stability otherwise were summarized as damaging stability; and variants which lost PTMs or regulatory motifs were summarized as damaging regulation. Fisher’s exact test was used to obtain odds ratios. To compare to the assay results of F. Mermet-Meillon et. al ^19^, we considered variants to be destabilizing by their assay when log_2_FC_sens_ ≤ - 0.3, and to impact cellular proliferation when |log_2_FC_prolif_| > 0.3.

## Acknowledgements

This research was completed in part with computational resources and technical support provided by the Research Computing Center at the Medical College of Wisconsin.

---

**Supplementary Figure 1:**
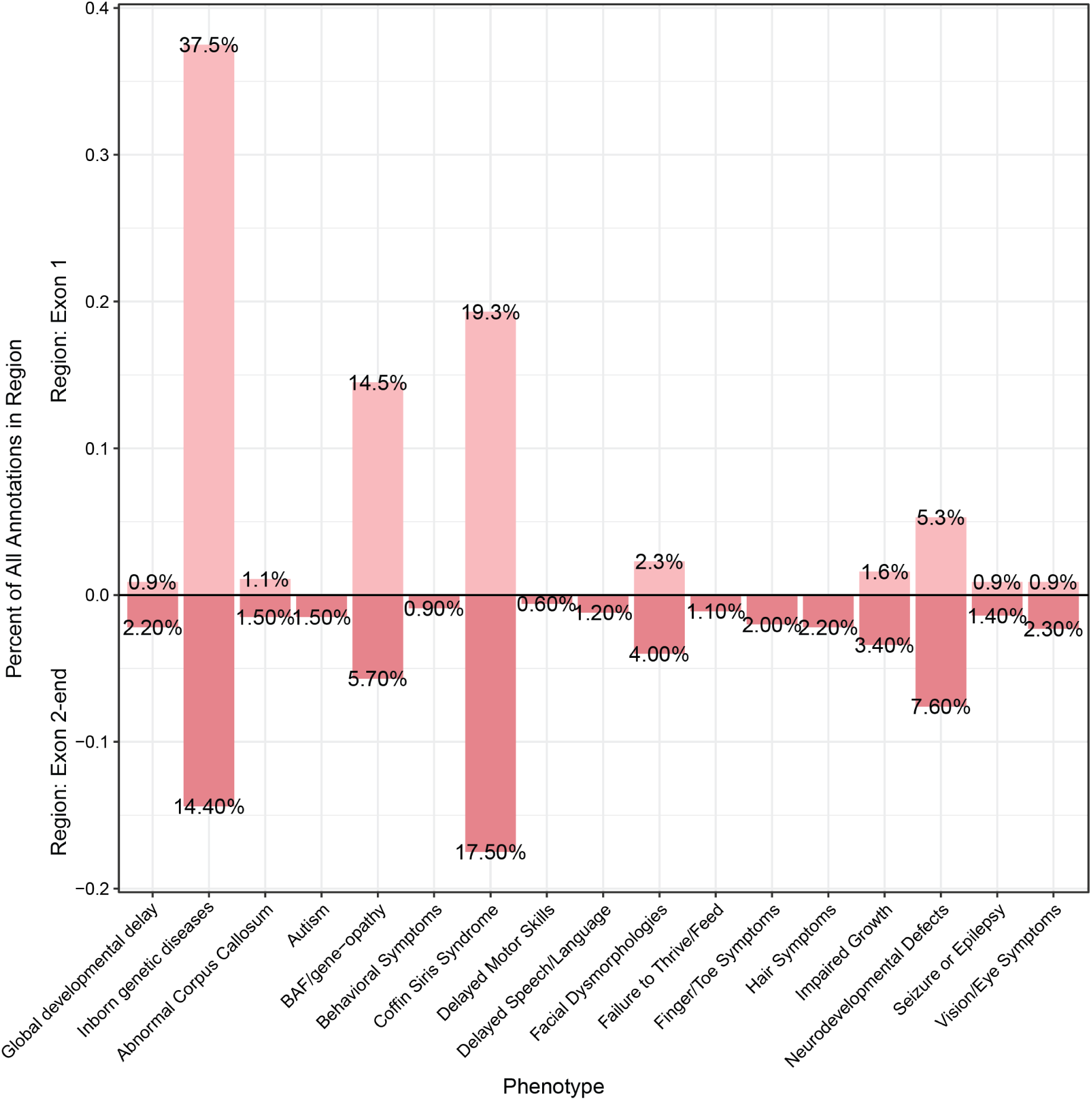
Variants in the first exon are associated with different traits than variants in the rest of the protein. The percent of all annotations in each region of the *ARID1B* gene is shown.

**Supplementary Figure 2:**
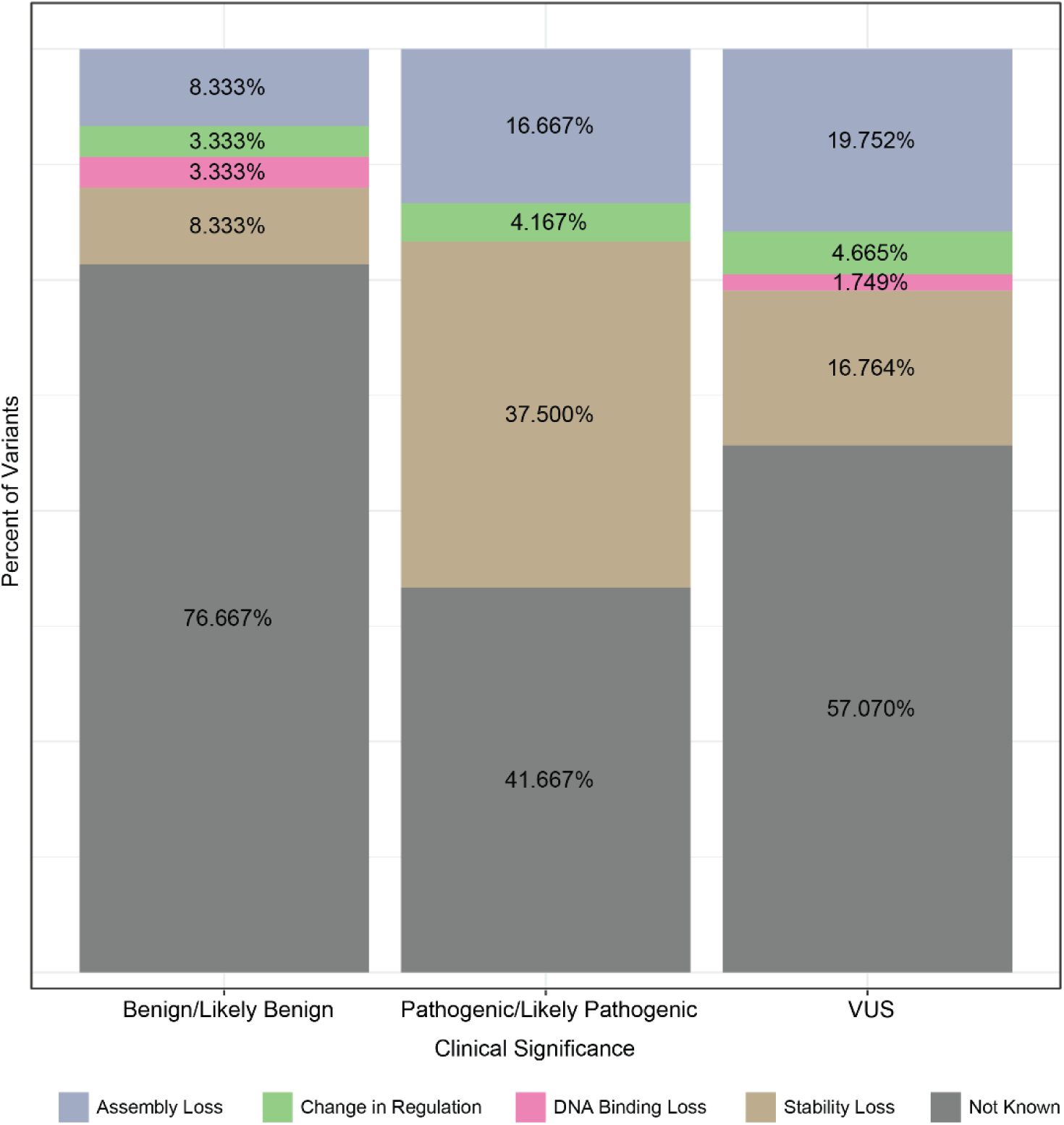
**Most Pathogenic variants cause stability loss, but many also cause loss of BAF assembly.**

**Supplementary Figure 3:**
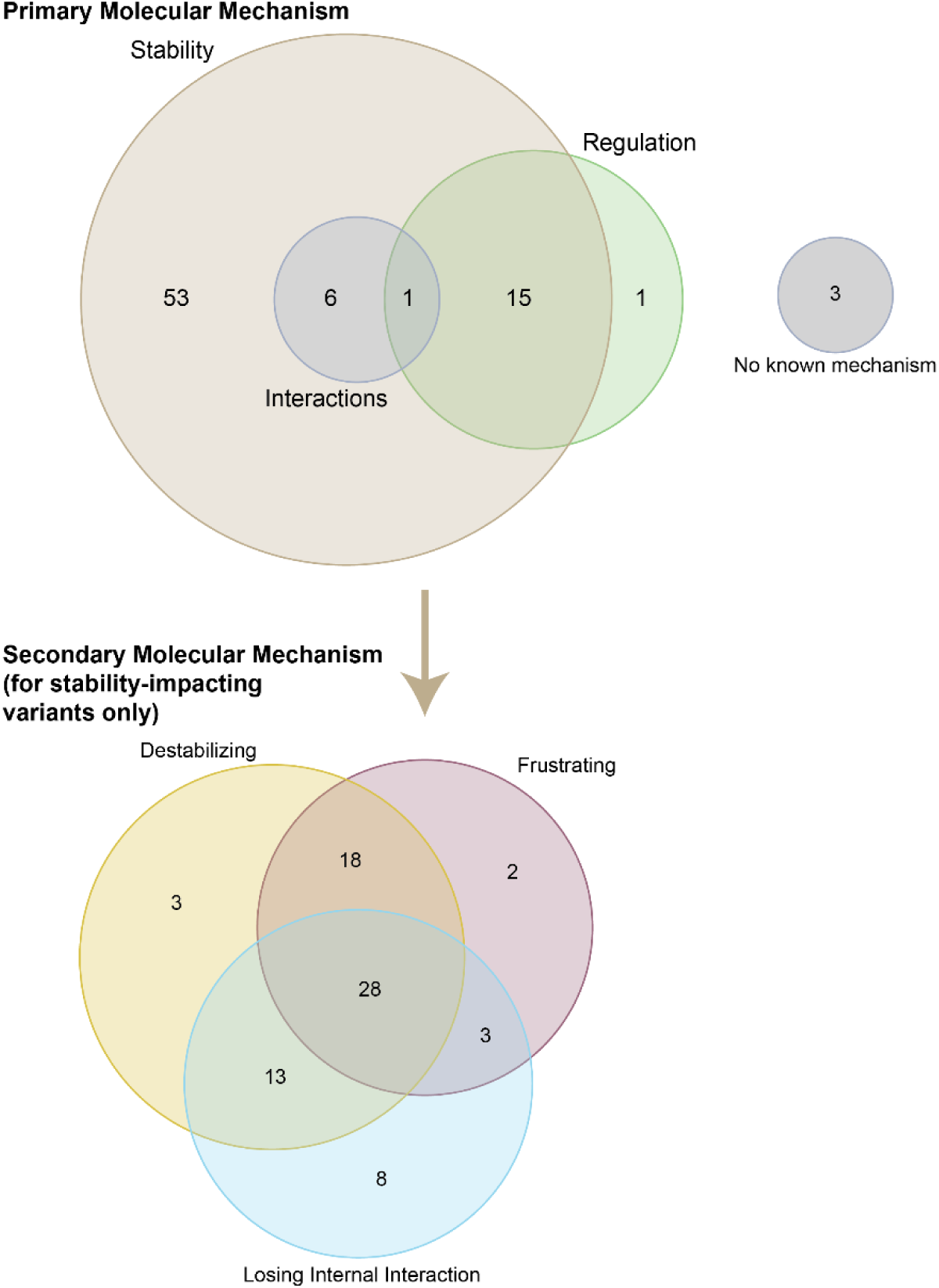
**Most abundance-loss proteins impact stability and regulation**. The primary molecular mechanisms impacting ARID1B abundance-loss variants is shown in the Euler diagram. The type of stability effect, as a secondary molecular mechanism, is shown in the second Euler diagram.

Supplementary Table 1: All missense variants and their annotations, calculations, and hypothesized effect on ARID1B functions. (Supplementary_Table1.xlsx)

## References

1. López-Rivera, J.A. et al. A catalogue of new incidence estimates of monogenic neurodevelopmental disorders caused by de novo variants. Brain 143, 1099–1105 (2020).

2. Gao, S., Shan, C., Zhang, R. & Wang, T. Genetic advances in neurodevelopmental disorders. Med Rev (2021) 5, 139–151 (2025).

3. Gillentine, M.A., Wang, T. & Eichler, E.E. Estimating the Prevalence of De Novo Monogenic Neurodevelopmental Disorders from Large Cohort Studies. Biomedicines 10, 2865 (2022).

4. Shain, A.H. & Pollack, J.R. The spectrum of SWI/SNF mutations, ubiquitous in human cancers. PLoS One 8, e55119 (2013).

5. Kadoch, C. et al. Proteomic and bioinformatic analysis of mammalian SWI/SNF complexes identifies extensive roles in human malignancy. Nat Genet 45, 592–601 (2013).

6. McRae, J.F. et al. Prevalence and architecture of de novo mutations in developmental disorders. Nature 542, 433–438 (2017).

7. Hein, K.Z., Stephen, B. & Fu, S. Therapeutic Role of Synthetic Lethality in ARID1A-Deficient Malignancies. J Immunother Precis Oncol 7, 41–52 (2024).

8. Helming, K.C. et al. ARID1B is a specific vulnerability in ARID1A-mutant cancers. Nat Med 20, 251–4 (2014).

9. Wang, Z. et al. Dual ARID1A/ARID1B loss leads to rapid carcinogenesis and disruptive redistribution of BAF complexes. Nat Cancer 1, 909–922 (2020).

10. Hellsvik, K. Foghorn Therapeutics Announces Updates for Selective ARID1B, Selective CBP and Selective EP300 Degrader Programs. (GLOBE NEWSWIRE, 2025).

11. Chen, K., Yuan, J., Sia, Y. & Chen, Z. Mechanism of action of the SWI/SNF family complexes. Nucleus 14, 2165604 (2023).

12. Egel, R., Beach, D.H. & Klar, A.J. Genes required for initiation and resolution steps of mating-type switching in fission yeast. Proceedings of the National Academy of Sciences 81, 3481–3485 (1984).

13. Kadoch, C. & Crabtree, G.R. Mammalian SWI/SNF chromatin remodeling complexes and cancer: Mechanistic insights gained from human genomics. Science Advances 1, e1500447 (2015).

14. Sandhya, S., Maulik, A., Giri, M. & Singh, M. Domain architecture of BAF250a reveals the ARID and ARM-repeat domains with implication in function and assembly of the BAF remodeling complex. PLoS One 13, e0205267 (2018).

15. He, S. et al. Structure of nucleosome-bound human BAF complex. Science 367, 875–881 (2020).

16. Hurlstone, A.F., Olave, I.A., Barker, N., van Noort, M. & Clevers, H. Cloning and characterization of hELD/OSA1, a novel BRG1 interacting protein. Biochem J 364, 255–64 (2002).

17. van der Sluijs, P.J. et al. ARID1B-related disorder in 87 adults: Natural history and self-sustainability. Genetics in Medicine Open 2(2024).

18. van der Sluijs, P.J. et al. A Case Series of Familial ARID1B Variants Illustrating Variable Expression and Suggestions to Update the ACMG Criteria. Genes (Basel) 12(2021).

19. Mermet-Meillon, F. et al. Protein destabilization underlies pathogenic missense mutations in ARID1B. Nature Structural & Molecular Biology 31, 1018–1022 (2024).

20. Chi, Y.I. et al. Structural bioinformatics enhances the interpretation of somatic mutations in KDM6A found in human cancers. Comput Struct Biotechnol J 20, 2200–2211 (2022).

21. Wagenknecht, J.B. et al. Multi-State Structural Genomics Enables Large-Scale, Mechanistic, and Context-Specific Classification of ABCC6 Genetic Variants Implicated in Calcification Diseases. Int J Mol Sci 27(2026).

22. Chen, S. et al. A genomic mutational constraint map using variation in 76,156 human genomes. Nature 625, 92–100 (2024).

23. Lo, T. et al. Sequencing of selected chromatin remodelling genes reveals increased burden of rare missense variants in ASD patients from the Japanese population. International Review of Psychiatry 34, 154–167 (2022).

24. Forwood, C. et al. Integration of EpiSign, facial phenotyping, and likelihood ratio interpretation of clinical abnormalities in the re-classification of an ARID1B missense variant. Am J Med Genet C Semin Med Genet 193, e32056 (2023).

25. Aspromonte, M.C. et al. Genetic variants and phenotypic data curated for the CAGI6 intellectual disability panel challenge. Hum Genet 144, 309–326 (2025).

26. Bosch, E. et al. The missing link: ARID1B non-truncating variants causing Coffin-Siris syndrome due to protein aggregation. Human Genetics 143, 965–978 (2024).

27. van der Sluijs, P.J. et al. The ARID1B spectrum in 143 patients: from nonsyndromic intellectual disability to Coffin–Siris syndrome. Genetics in Medicine 21, 1295–1307 (2019).

28. Sim, J.C., White, S.M. & Lockhart, P.J. ARID1B-mediated disorders: Mutations and possible mechanisms. Intractable Rare Dis Res 4, 17–23 (2015).

29. Khalil, A., Collins, M.P. & Quintás-Cardama, A. Prognostic Implications of SMARCA4, ARID1A, and Other BAF Mutations in Non-Small Cell Lung Cancer. Cancer Med 14, e71442 (2025).

30. Bayraktar, R. et al. The mutational landscape and functional effects of noncoding ultraconserved elements in human cancers. Science Advances 11, eado2830 (2025).

31. Patil, A. et al. A disordered region controls cBAF activity via condensation and partner recruitment. Cell 186, 4936–4955.e26 (2023).

32. Gerasimavicius, L., Teichmann, S.A. & Marsh, J.A. Leveraging protein structural information to improve variant effect prediction. Curr Opin Struct Biol 92, 103023 (2025).

33. Tripathi, S., Dsouza, N.R., Mathison, A.J., Leverence, E., Urrutia, R. & Zimmermann, M.T. Enhanced interpretation of 935 hotspot and non-hotspot RAS variants using evidence-based structural bioinformatics. Comput Struct Biotechnol J 20, 117–127 (2022).

34. Dsouza, N.R. et al. Structural and Dynamic Analyses of Pathogenic Variants in PIK3R1 Reveal a Shared Mechanism Associated among Cancer, Undergrowth, and Overgrowth Syndromes. Life (Basel) 14(2024).

35. Deng, K. et al. Molecular mechanisms of TTC21B gene mutations in nephronophthisis type 12 and genetic prevention through PGT. Frontiers in Genetics Volume 16 - 2025(2025).

36. Landrum, M.J. et al. ClinVar: improvements to accessing data. Nucleic Acids Res 48, D835–D844 (2020).

37. Fokkema, I.F.A.C. et al. The LOVD3 platform: efficient genome-wide sharing of genetic variants. European Journal of Human Genetics 29, 1796–1803 (2021).

38. Bick, A.G. et al. Genomic data in the All of Us Research Program. Nature 627, 340–346 (2024).

39. Tate, J.G. et al. COSMIC: the Catalogue Of Somatic Mutations In Cancer. Nucleic Acids Res 47, D941–D947 (2019).

40. Gao, J. et al. Integrative analysis of complex cancer genomics and clinical profiles using the cBioPortal. Sci Signal 6, pl1 (2013).

41. Dyer, S.C., et al. Ensembl 2025. Nucleic Acids Research 53, D948–D957 (2025).

42. Siegel, R.L., Kratzer, T.B., Giaquinto, A.N., Sung, H. & Jemal, A. Cancer statistics, 2025. CA: A Cancer Journal for Clinicians 75, 10–45 (2025).

43. Necci, M., Piovesan, D., Dosztanyi, Z. & Tosatto, S.C.E. MobiDB-lite: fast and highly specific consensus prediction of intrinsic disorder in proteins. Bioinformatics 33, 1402–1404 (2017).

44. Kumar, M. et al. ELM—the eukaryotic linear motif resource in 2020. Nucleic Acids Research 48, D296–D306 (2019).

45. Hornbeck, P.V., Zhang, B., Murray, B., Kornhauser, J.M., Latham, V. & Skrzypek, E. PhosphoSitePlus, 2014: mutations, PTMs and recalibrations. Nucleic acids research 43, D512–20 (2015).

46. Pommié, C., Levadoux, S., Sabatier, R., Lefranc, G. & Lefranc, M.P. IMGT standardized criteria for statistical analysis of immunoglobulin V-REGION amino acid properties. J Mol Recognit 17, 17–32 (2004).

47. Abramson, J. et al. Accurate structure prediction of biomolecular interactions with AlphaFold 3. Nature 630, 493–500 (2024).

48. H. Niwa, A.S., S. Yokoyama. Crystal structure of the HBAF250B at-rich interaction domain (ARID). (RIKEN Structural Genomics/Proteomics Initiative (RSGI), 2007).

49. He, S., Wu, Z., Tian, Y., Yu, Z., Yu, J., Wang, X., Li, J., Liu, B., Xu, Y. Structure of nucleosome-bound human BAF complex. (2020).

50. Schake, P., Bolz, S.N., Linnemann, K. & Schroeder, M. PLIP 2025: introducing protein–protein interactions to the protein–ligand interaction profiler. Nucleic Acids Research 53, W463–W465 (2025).

51. Grant, B.J., Rodrigues, A.P., ElSawy, K.M., McCammon, J.A. & Caves, L.S. Bio3d: an R package for the comparative analysis of protein structures. Bioinformatics 22, 2695–6 (2006).

52. Schymkowitz, J., Borg, J., Stricher, F., Nys, R., Rousseau, F. & Serrano, L. The FoldX web server: an online force field. Nucleic acids research 33, W382–8 (2005).

53. Parra, R.G. et al. Protein Frustratometer 2: a tool to localize energetic frustration in protein molecules, now with electrostatics. Nucleic Acids Res 44, W356–60 (2016).

